# Abiotic Stress-Mediated Modulation of the tncRNome: Unraveling tRNA-Derived RNA Networks in Plant Adaptive Responses

**DOI:** 10.64898/2025.12.04.692324

**Authors:** Shafaque Zahra, Rashmi Gangwar, Sneha Tiwari, Dipul Kumar Biswas, Amarjeet Singh, Shailesh Kumar

## Abstract

Transfer RNA-derived non-coding RNAs (tncRNAs) are potential regulators of gene expression in plants, and it’s crucial to understand their diversity and functional roles in modulating plant stress responses. Here, we have investigated abiotic stress-responsive tncRNAs in two important crops, rice and chickpea, under drought and salt stress conditions. Small RNA sequencing revealed that 70.2% of tncRNAs in rice and 45% in chickpea were commonly expressed across multiple stress conditions. qRT-PCR validation showed that most tRF-5s were upregulated in both rice and chickpea under drought and salt stress. In rice, 3′tRHs were specifically induced by drought, whereas 5′tRHs responded predominantly to salt stress. In chickpea, tRF-3s were upregulated during drought but downregulated under salt stress.

Furthermore, we confirmed the mRNA targets of these stress-associated tRNA fragments using RLM-RACE, thereby verifying the cleavage of specific target mRNAs in both species. Expression profiling of target genes revealed a negative association with their corresponding tRFs in both species. Subcellular localization demonstrated that OsGEM and OsPPR were primarily localized in the nucleus, while CaMIZ1 and CaSKP1 were detected at the plasma membrane. Overall, we identified abiotic stress-responsive tncRNAs from rice and chickpea that suggest their potential contribution to plant resilience under changing environmental conditions.

## Introduction

The tRNA-derived non-coding RNAs or tncRNAs are generated through the precise enzymatic cleavage of tRNAs into smaller fragments under specific conditions, such as abiotic and biotic stress, developmental transitions, nutrient deprivation, or pathogenic attacks, where they play regulatory roles in gene expression and stress responses. The tncRNA includes 12-30 nt length tRFs (transfer RNA-derived fragments), and longer tRNA halves or tRHs (∼30 to 40 nt). Based on their cleavage site on mature tRNA and length, they are further classified into various classes, including tRF-5, tRF-3, i-tRF, 5’tRNA half/5’tRH, and 3’tRNA half/3’tRH. Precursor tRNAs also give rise to lesser-studied tRF-1, leader tRF, and other tRFs. Eukaryotic tRFs and tRHs are produced by precise cleavage of tRNA through a Dicer-dependent or -independent pathway (Kuscu et al., 2018). In plants, they are generated through a DCL (Dicer-like)-independent pathway, e.g., the ribonucleases RNS1-mediated process (Alves and Nogueira, 2021). Biogenesis of tRFs is a non-random process and can mimic the microRNA generation pathway (Venkatesh, Suresh, and Tsutsumi, 2016). The existence of tRFs, their diverse classes, and their largely unexplored functions in plants necessitate further research on tRNA beyond its role as an adapter molecule.

The existence of tRFs in plants was first reported in *Arabidopsis* (Thompson et al., 2008). Just as with microRNAs (miRNAs), some of these fragments are associated with proteins such as Argonaute (AGO). These proteins are crucial players in RNA silencing machinery and function by forming RISC (RNA-induced silencing complex) (Kumar et al., 2014; Kuscu et al., 2018). Some reports show the differential expression of tRFs in response to biotic and abiotic stress responses in plants (Alves and Nogueira, 2021). For instance, fragments derived from AspGTC and GlyTCC tRNAs were expressed in abundance during phosphate starvation and drought stress in *Arabidopsis* (Hsieh et al., 2009). In 2013, it was reported that under drought stress, ∼19 nt tRFs derived from AlaAGC, ArgCCT, ArgTCG, and GlyTCC tRNAs were expressed in abundance in *Arabidopsis* (Loss-Morais et al., 2013). These fragments were also expressed in response to cold, whereas tRFs from ArgTCG and TyrGTA were generated during oxidative stress(Alves et al., 2017).

tncRNAs exhibit remarkable diversity and versatility in regulating various biological processes (Fig. 1a). However, their expression patterns, biogenesis, and specific cellular functions in plants remain insufficiently understood. Comprehensive studies on these aspects are essential to clarify how tncRNAs contribute to adaptive responses under abiotic stresses, such as drought and salinity, which are major factors severely limiting crop productivity. While earlier studies on tncRNAs have focused on model plants like *Arabidopsis*, their roles in major crops remain largely unexplored. This study bridges that gap by providing the first comprehensive identification, experimental validation, and functional analysis of stress-associated tncRNAs in rice and chickpea. Some databases of tncRNAs, such as PtRFdb (Gupta et al., 2018), tRex (Thompson et al., 2018), and PtncRNAdb (Zahra et al., 2022), and a toolkit, i.e., tncRNA-toolkit (Zahra et al., 2021 and Zahra et al., 2023), have also been developed for the identification of tncRNAs. Recently, our research group identified and annotated different classes of tncRNA in small RNA sequencing datasets across six plant species (Zahra et al., 2021).

**Figure 1:**
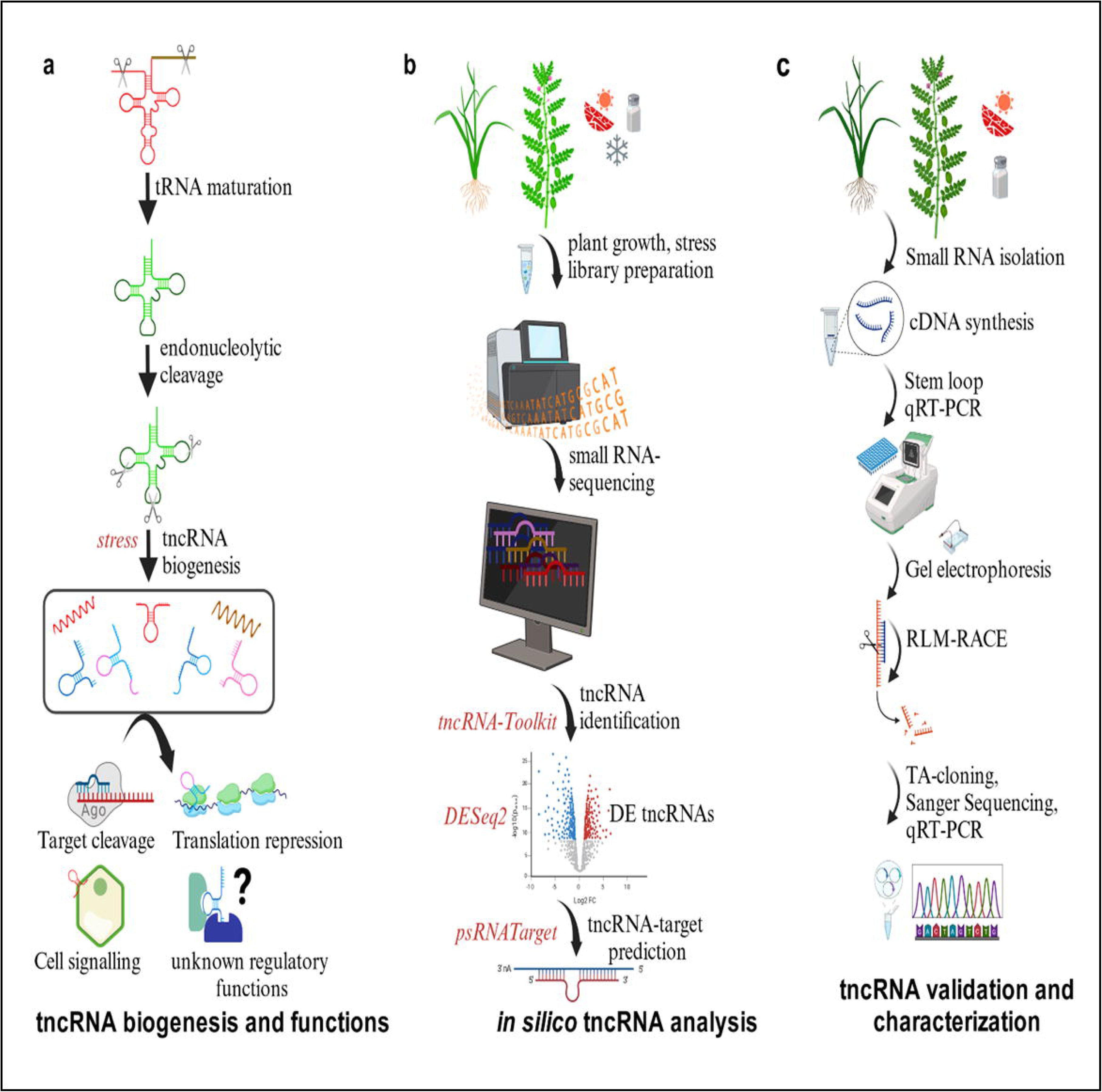
tncRNA generation, functions, as well as their identification workflow applied in this study. (a) Biogenesis of tncRNAs from pre- and mature tRNAs based on various cleavage sites on their progenitor tRNA molecules, as well as biological functions in the cellular milieu. (b) Sequential workflow of high throughput *in silico* tncRNA identification in response to abiotic stress conditions in Rice and Chickpea. (c) Experimental validation of potential differentially expressed (DE) tncRNAs and their putative targets in Rice and Chickpea (Created with BioRender.com).

Building upon this foundation, the current work significantly advances the understanding of tncRNAs by leveraging comprehensive in-house small RNA sequencing datasets from two important crop species, rice (*Oryza sativa*) and chickpea (*Cicer arietinum*), under abiotic stress conditions. It’s an integrated approach combining computational prediction with experimental validation to identify and characterize distinct classes of tncRNAs and their specific target genes (Figs 1b, c). Through small RNA sequencing of rice and chickpea seedlings, we identified 5,282 unique tncRNAs in rice and 1,630 in chickpea under drought, while under salt stress, 4,470 were found in rice and 1,875 in chickpea (Table 1). A subset of these tncRNAs: 18 from rice and 31 from chickpea (Supplementary Tables S1.1, S1.2) were further screened for their expression profiles and experimentally validated, resulting in the identification of 14 rice and 26 chickpea tncRNAs that are differentially expressed in response to stress conditions (Tables 2, 3). Moreover, we have demonstrated that certain tncRNAs possess the ability to precisely target and cleave specific mRNAs, thereby modulating gene expression under adverse environmental conditions. To substantiate these findings, we conducted *in silico* predictions of tncRNA-mRNA interactions followed by rigorous experimental validation using RNA ligase-mediated rapid amplification of cDNA ends (RLM-RACE) experiments. These insights expand our understanding of the molecular mechanisms underlying plant stress resilience and pave the way for future research into the functional roles of tncRNAs in crop improvement strategies.

**Table 1:** Summary of identified groups of transfer RNA-derived non-coding RNAs (tncRNAs) abundance i Rice and Chickpea under control and abiotic stress conditions.

| Plant Species | Conditions | tncRNAs | tRF-5 | 5'tRH | tRF-3(CCA) | 3'tRH(CCA) | tRF-1 | Leader tRF | Other tRFs |
| --- | --- | --- | --- | --- | --- | --- | --- | --- | --- |
|  |  | Total/Unique | Total/Unique | Total/Unique | Total/Unique | Total/Unique | Total/Unique | Total/Unique | Total/Unique |
| Rice<br>(*Oryza sativa*) | Control | 13351/5871 | 1922/753 | 431/197 | 2820/1131 | 13/11 | 28/18 | 2/1 | 8137/3760 |
|  | Drought | 8855/5282 | 1598/856 | 517/260 | 1941/1051 | 37/27 | 19/13 | 1/1 | 4742/3074 |
|  | Salt | 10016/4470 | 1651/673 | 596/248 | 2200/916 | 48/25 | 39/23 | 0/0 | 5482/2585 |
| Chickpea<br>(*Cicer arietinum*) | Control<br>_Drought | 1803/885 | 396/193 | 163/73 | 808/368 | 78/34 | 11/5 | 4/2 | 347/210 |
|  | Drought | 3042/1630 | 652/305 | 253/114 | 1384/705 | 122/55 | 13/5 | 6/2 | 612/444 |
|  | Control<br>_Salt | 2577/1254 | 539/238 | 215/98 | 1082/463 | 99/38 | 11/5 | 6/4 | 625/408 |
|  | Salt | 3464/1875 | 654/349 | 227/109 | 1616/712 | 83/40 | 7/7 | 3/3 | 874/655 |

**Table 2:**
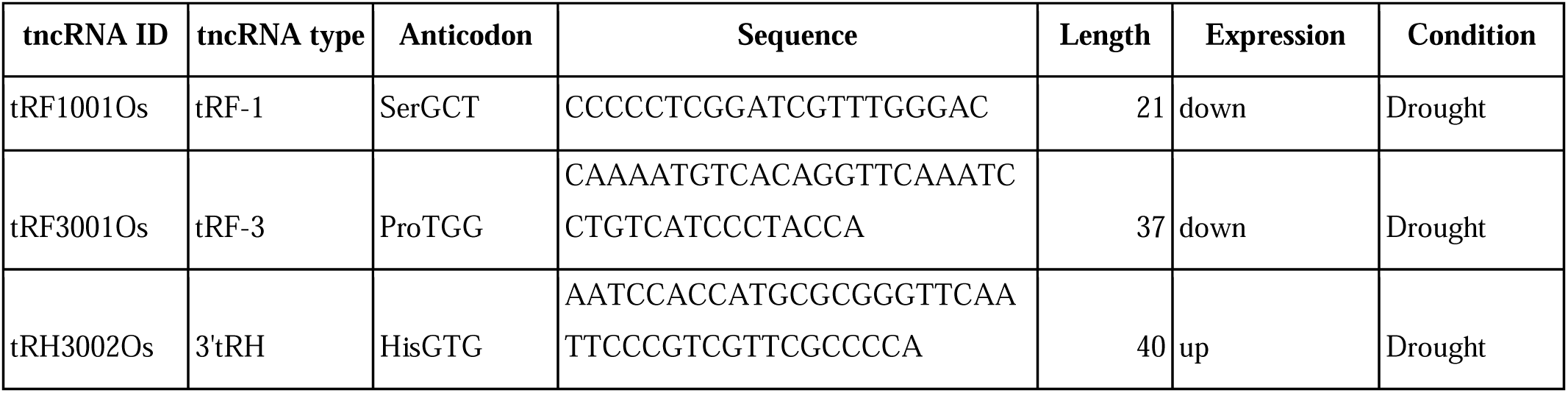
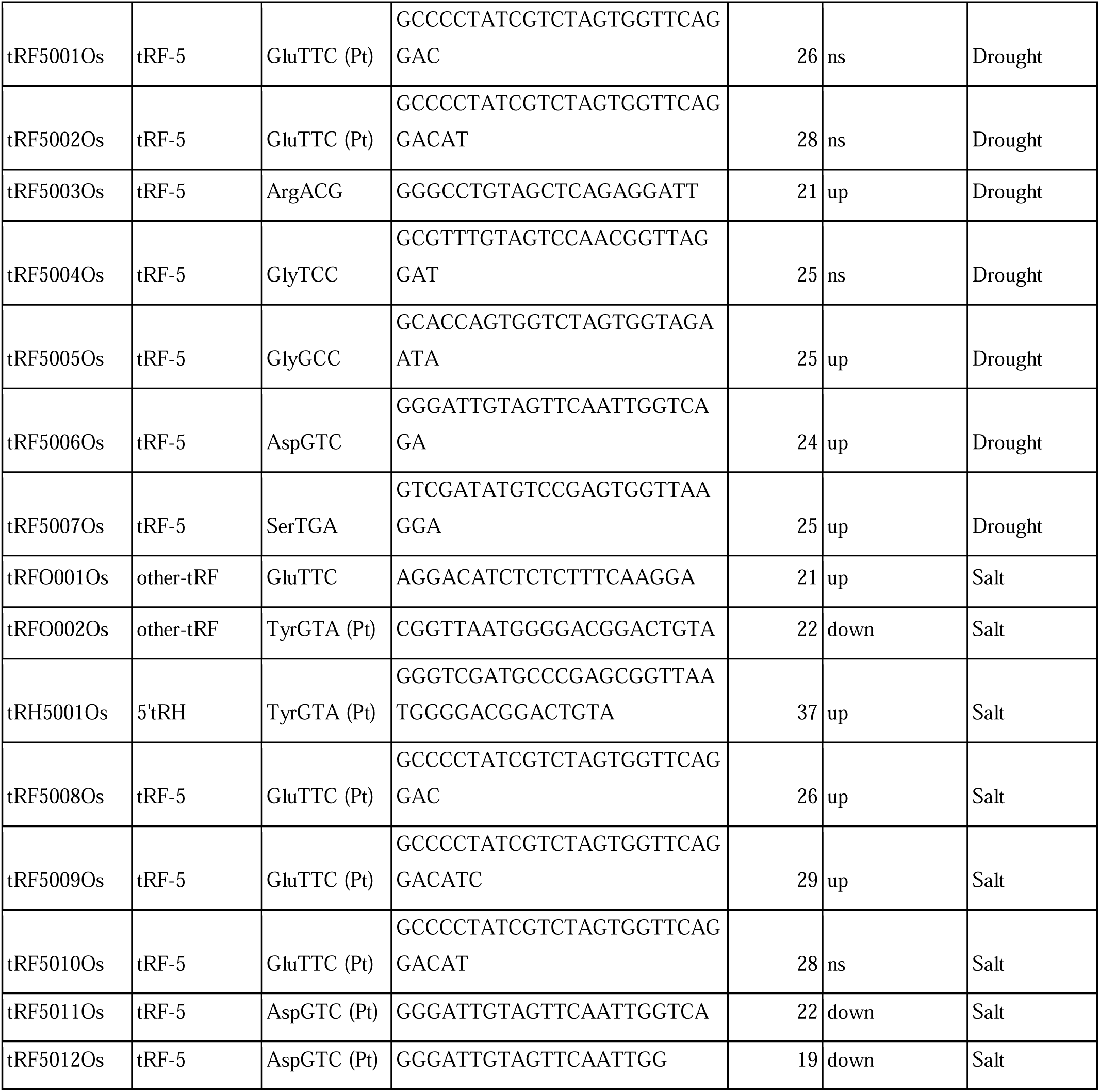
Experimentally validated tncRNAs along with other details related to their type, tRNA origin selected for Rice datasets (up: significantly upregulated, down: significantly downregulated, ns: not significantly dysregulated; Pt: plastidial).

**Table 3:**
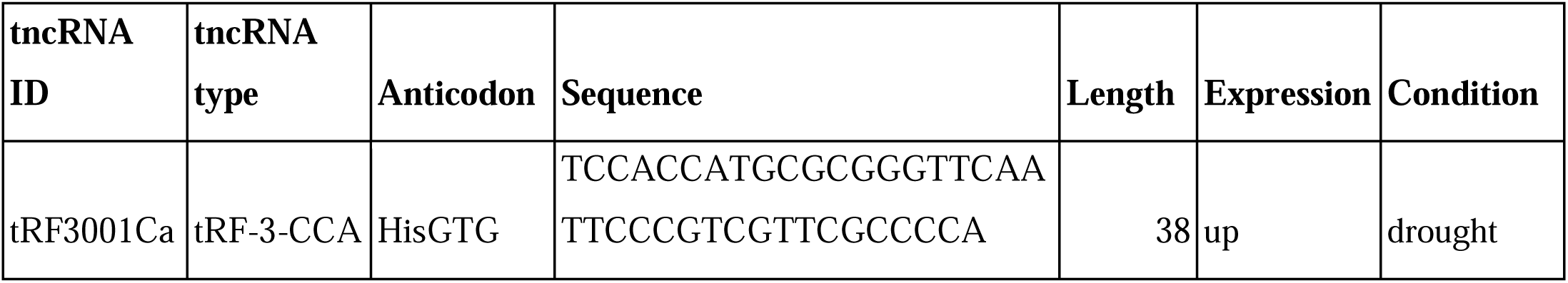
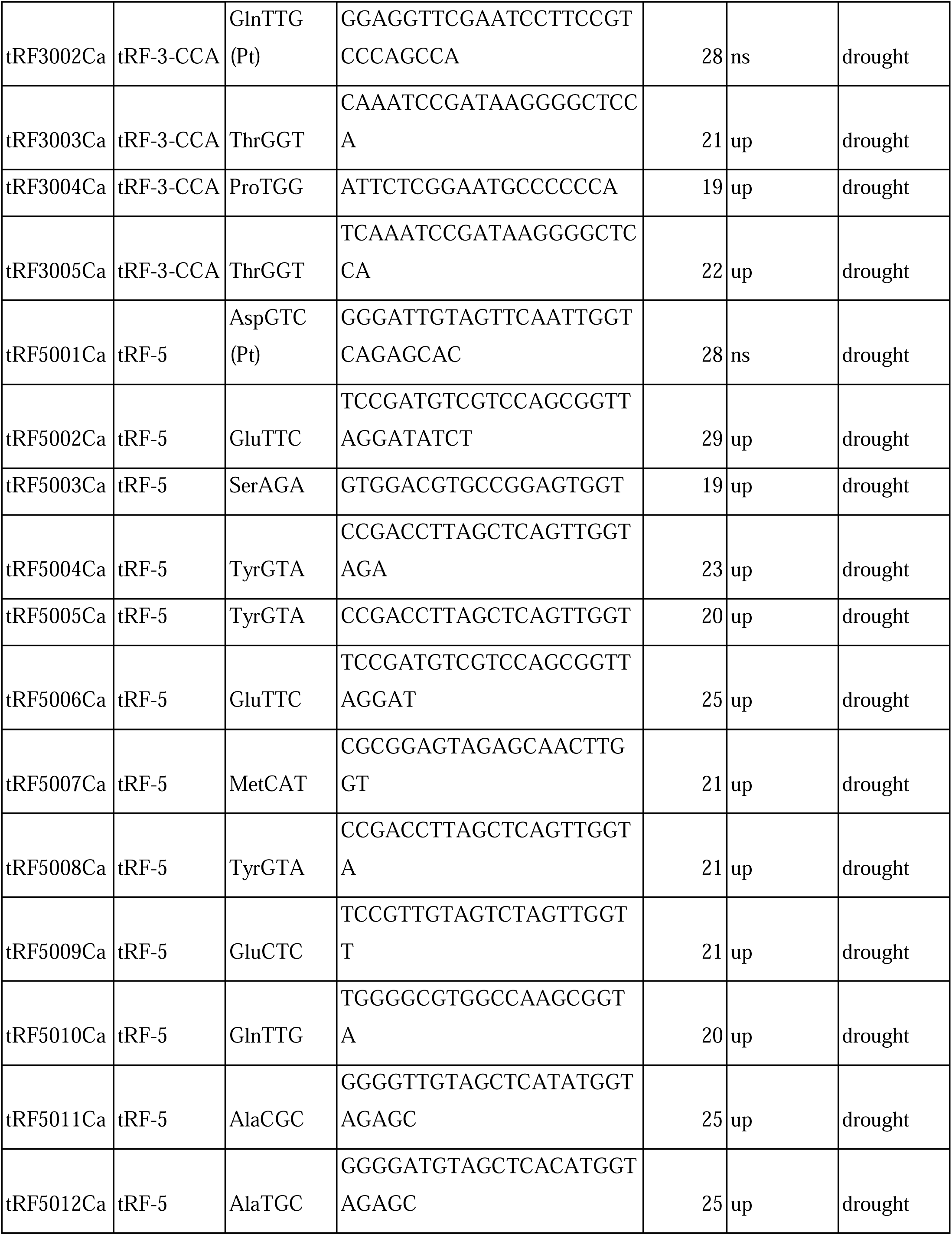
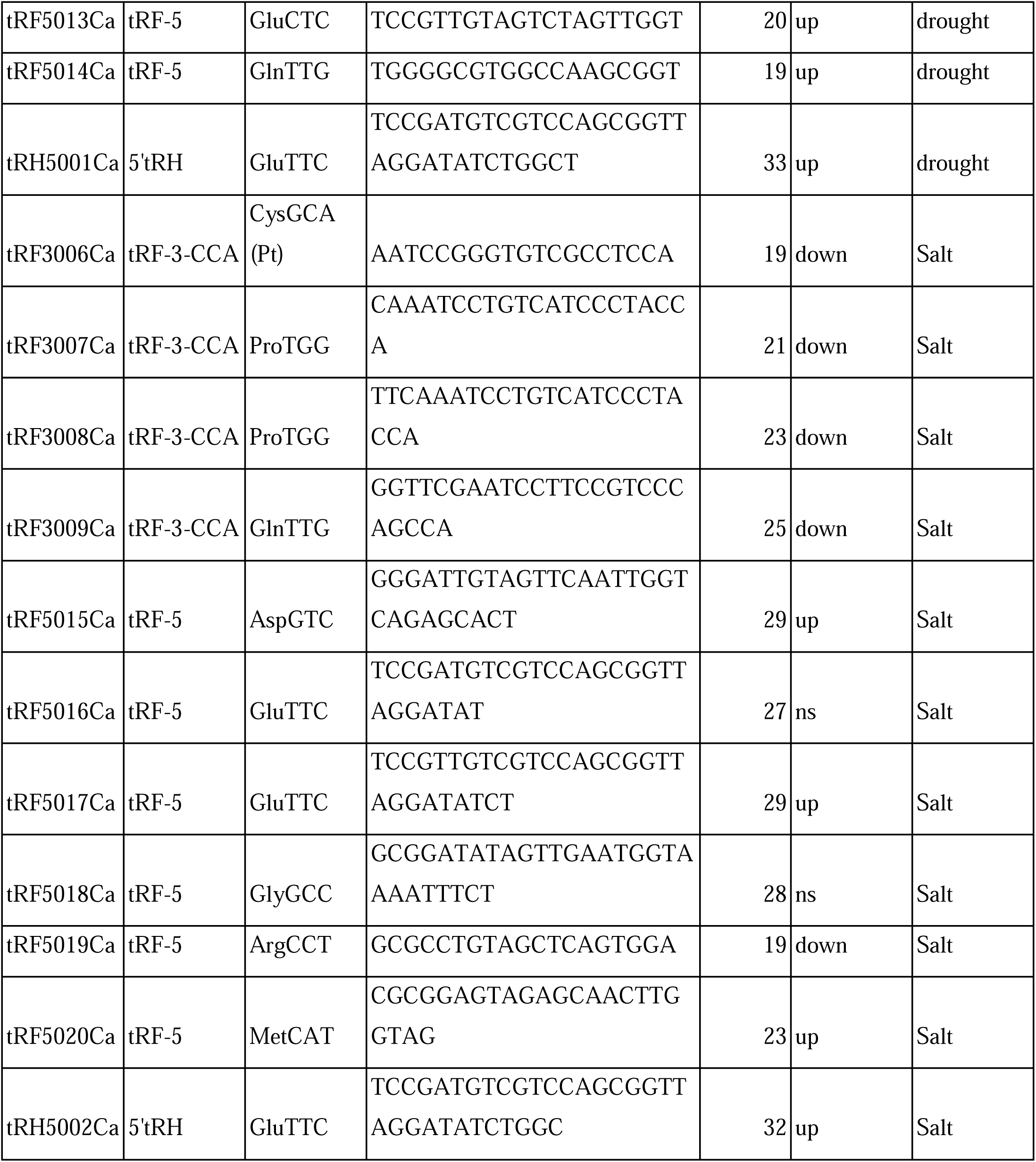
Experimentally validated tncRNAs along with other details related to their type, tRNA origin selected for Chickpea datasets (up: significantly upregulated, down: significantly downregulated, ns: not significantly dysregulated; Pt: plastidial).

## Materials and Methods

### Plant materials, growth conditions, and abiotic stress treatments

*Oryza sativa* ssp. *indica* (rice) genotype IR64, and *Cicer arietinum* L. (chickpea) genotype ICC4958 were used for stress experiments. Rice seeds were surface-sterilized, soaked in distilled water for 24 hours at 37°C, and germinated for one week on MS agar medium. The coleoptile of germinated seeds was transferred to Styrofoam supports and placed in Yoshida nutrient solution and grown in a controlled growth chamber with a 16 h/8 h light/dark photoperiod at 26°C/28°C (day/night) temperatures under a light intensity of 200 μmol m□² s□¹. Chickpea seeds were grown in an autoclaved 1:1 mixture of agro-peat and vermiculite in plastic pots and maintained at 22°C under a 14-hour photoperiod with a light intensity of approximately 250 µmol m□² s□¹ (Garg et al., 2010).

The 21-day-old seedlings of both species were used for abiotic stress treatment. For drought stress, rice seedlings were exposed to a 20% (w/v) solution of polyethylene glycol (PEG) 6000, while chickpea seedlings were placed on tissue paper folds for 5 hours to impose rapid dehydration (Garg et al., 2010). Chickpea and rice seedlings were exposed to a 150 mM NaCl solution for salinity stress. For both species, control seedlings were maintained under normal growth conditions. Following 5 hours of treatment, tissue samples were collected from both stressed and control plants. For each condition, three separate biological replicates were harvested, immediately frozen in liquid nitrogen, and stored at -80°C until further use.

### Small RNA extraction and library preparation

Small RNAs were subsequently isolated from control and stressed seedling tissues of rice and chickpea using the High Pure miRNA Isolation Kit (Merck, Germany) and visualized on a 15% urea denaturing polyacrylamide gel (PAGE). The concentration of small RNA samples was measured using a NanoDrop 1000 spectrophotometer (Thermo Fisher Scientific, USA). For small RNA library preparation, after quality assessment using the Agilent 2100 Bioanalyzer, 1 µg of high-quality small RNA samples (in three biological replicates per condition) from rice and chickpea were submitted for library preparation and sequencing using single-end 50 bp reads on the Illumina HiSeq 2500 platform, generating approximately 15–20 million reads per library.

### Small RNA-sequencing data processing and tncRNA identification

We carried out an analysis of single-end small RNA-Seq datasets (Illumina) generated from rice and chickpea. The raw single-end reads were processed using the fastp tool at default settings (Chen et al., 2018). Genome assembly for *Cicer arietinum* (nuclear: ASM33114v1; organellar: ASM33114v1) and *Oryza sativa* (Build 4.0; organellar: IRGSP-1.0) was utilized for the identification of tRNAs. Identification and annotation of tncRNAs were performed using tncRNA-Toolkit as done in our previous study (Zahra et al., 2023).

### Differentially expressed tncRNAs detection

Differentially expressed tncRNAs were identified using DESeq2 (Love et al., 2014), as previously done in our earlier study (Zahra et al., 2021). The tncRNAs with a p-value less than 0.05 were considered significant. Those tncRNAs, with log2FC value greater than or equal to 1 (≥□1) and less than or equal to -1 (≤□-1), were considered up- and down-regulated, respectively.

### Stem-Loop RT-PCR and qPCR Validation of tncRNAs Under Abiotic Stress

For tncRNAs expression analysis, tRF-specific stem-loop primers were designed with forward primers corresponding to the full mature tRF sequences and a universal reverse primer (Supplementary Table S2). For stem-loop RT-PCR, 1 μg of small RNA was hybridized with tRF-specific stem-loop forward primers, and the reaction mixture included 1× RT buffer, 0.5 mM dNTPs, and 200 U of M-MLV reverse transcriptase. Reverse transcription was carried out at 16°C for 30 minutes, followed by 42°C for 60 minutes, and terminated at 70°C for 5 minutes. The expression patterns of tRFs under various abiotic stress conditions (drought and salinity) were evaluated using SYBR Green I Master Mix (Applied Biosystems) by Applied Biosystems 7900 Real-Time PCR System. For normalization, 18S and 25S rRNA genes were used as endogenous controls (Garg et al., 2010; Jain et al., 2006). Relative fold change was calculated using the 2^-ΔΔCt^ method (Livak and Schmittgen, 2001). Each data point represents the mean of three biological replicates, each with three technical replicates. Graphs were prepared in GraphPad Prism 8.0. Statistical significance was assessed using Student’s *t*-test.

### Prediction and Functional Enrichment of Target Genes for Validated tncRNAs

The putative target genes of validated rice and chickpea tncRNAs (19–25 nt) under drought and salt stress were predicted using the psRNATarget server (2017 release, default parameters) (Dai et al., 2018). For rice, the *Oryza sativa* transcript, RAP-DB, v1, was used, and for chickpea, the *Cicer arietinum* transcript, ASM33114v1, was used as the reference database. To visualize potential regulatory interactions, ten predicted target genes, with priority given to those functionally linked to stress-responsive pathways, were selected. The interaction networks between tRFs and their predicted targets were constructed using Cytoscape (v3.10.4).

Functional enrichment analysis of drought- and salt-associated target genes in rice and chickpea was performed using the Database for Annotation, Visualization and Integrated Discovery (DAVID, version 6.8) (Sherman et al., 2022). GO analysis in three categories: biological processes (BP), cellular components (CC), and molecular functions (MF), along with KEGG pathway analysis, was performed (p < 0.05 was considered significant) (Huang et al., 2009).

### RLM-RACE and qRT-PCR-Based Validation of Predicted tRF Targets

Target mRNA cleavage sites were experimentally mapped by RNA ligase-mediated rapid amplification of cDNA ends (5′ RLM-RACE). To examine tRF-directed cleavage of predicted targets in vivo, total RNA was extracted from rice and chickpea seedlings using TRI Reagent (Sigma, USA), and a modified procedure for 5′ RLM-RACE was followed using the FirstChoice RLM-RACE Kit® (ThermoFisher Scientific/Invitrogen, Carlsbad, CA, USA) to amplify the 5′ UTR region of target gene mRNAs, according to the manufacturer’s protocol. Total RNA was ligated to the 5′ RNA adapter without tobacco acid pyrophosphatase (TAP) treatment to specifically detect tRF-mediated cleavage products. The Primer combinations used in the RLM-RACE experiments are listed in Supplementary Table S2. PCR products were visualized on agarose gel, purified using a Zymoclean Gel DNA Recovery Kit (ZYMO RESEARCH), cloned into the pCR4-TOPO TA vector (ThermoFisher Scientific), and sequenced for further analysis. For expression analysis of target genes under abiotic stress conditions, total RNA was extracted from the same rice and chickpea samples used for small RNA sequencing using TRI Reagent (Sigma, USA). Residual genomic DNA was removed by DNase I treatment (Thermo Scientific, USA), and RNA integrity was assessed by electrophoresis on a 1.5% denaturing agarose gel. Gene-specific primers were designed for qRT-PCR (Supplementary Table S2). First-strand cDNA was synthesized from 2 μg of total RNA using the Verso cDNA Synthesis Kit (Thermo Fisher Scientific). qRT-PCR was performed on an Applied Biosystems 7900 Real-Time PCR System using SYBR Green I Master Mix (Applied Biosystems). Expression levels were normalized to the endogenous control gene Elongation factor 1-alpha (EF-1α) (Garg et al., 2010; Jain et al., 2006) and relative fold changes were calculated using the 2^-ΔΔCt^ method (Livak and Schmittgen, 2001). Each data point represents the average of three biological and three technical replicates. Statistical significance was assessed using Student’s t-test.

### Subcellular localization of Target genes

The full-length coding sequences of *OsGEM* (XM_015776041.2, 744 bp), *OsPPR* (XM_015780543.2, 1173 bp) from *Oryza sativa* ssp. *indica* (rice) genotype IR64, and *CaSKP1* (XM_004515068.3, 1158 bp), *CaMIZ1* (XM_004515889.3, 729 bp) from *Cicer arietinum* L. (chickpea) genotype ICC4958 were PCR amplified using gene-specific primers (Supplementary Table S2) and cloned in-frame with a GFP tag into the pCAMBIA1300 vector. Positive clones were confirmed by PCR, restriction digestion, and sequencing. Verified clones were then transformed into *Agrobacterium tumefaciens* strain GV3101 for transient expression in *Nicotiana benthamiana*. *Agrobacterium* cultures carrying the GFP-tagged constructs or the empty GFP vector (control) were grown and adjusted to an optical density (OD□□□ ≈ 0.5) before infiltration. Cultures expressing either a plasma membrane marker (PM-RFP) or a nuclear marker (NM-RFP) were co-infiltrated into *N. benthamiana* leaves at a 1:1 ratio. After 48 hours, confocal microscopy was performed using a Leica TCS SP8 confocal laser-scanning microscope (Leica Microsystems, Germany) to capture fluorescent signals. The excitation and emission wavelengths were set at 488 nm and 500–540 nm for GFP, and at 561 nm and 580–620 nm for RFP, respectively.

### Statistical analysis

All experimental data were expressed as the mean with standard deviation (mean ± SD) of three independent biological replicates. Student’s t-test was performed to reveal significant differences between control and stress samples within each tncRNA class. One-way ANOVA was performed to compare tncRNA classes under both control and stress conditions using GraphPad Prism version 8 (Supplementary Table S3).

## Results

### Identification of tncRNAs in rice and chickpea under abiotic stress

Using our in-house small RNA-seq data, we identified a diverse set of tncRNAs in rice and chickpea under abiotic stress conditions. These tncRNA classes were categorized into tRF-5, tRF-3, tRF-1, leader-tRF, 5′tRH, 3′tRH, and other-tRFs, following the classification used in our previous study (Zahra et al., 2021). In rice, a total of 8855 tncRNAs were detected under drought and 10016 under salt stress, while in chickpea, 3042 and 3464 tncRNAs were identified under drought and salt conditions, respectively (Table 1). After an initial filtration based on unique sequences of tncRNAs class, a final unique set of each tncRNA class was obtained for both rice and chickpea across different conditions. The category-wise distribution of unique counts and normalized read counts of tncRNA fragments across different conditions is shown in Figure 2 (a–d). In both rice and chickpea, the tRF-5, tRF-3, and other-tRF classes consistently had a higher number of unique sequences compared to the remaining classes (Figs 2a, b). In contrast, the read counts for the other-tRF class were significantly lower than those of tRF-5 and tRF-3 under all conditions in both crops (Figs 2c, d). Interestingly, in rice, tRF-5 showed significantly higher unique sequences and other-tRF showed lower under salt stress compared to control (Fig. 2a), whereas in chickpea, tRF-3 showed a significant increase under salt stress relative to control (Fig. 2b).

**Figure 2:**
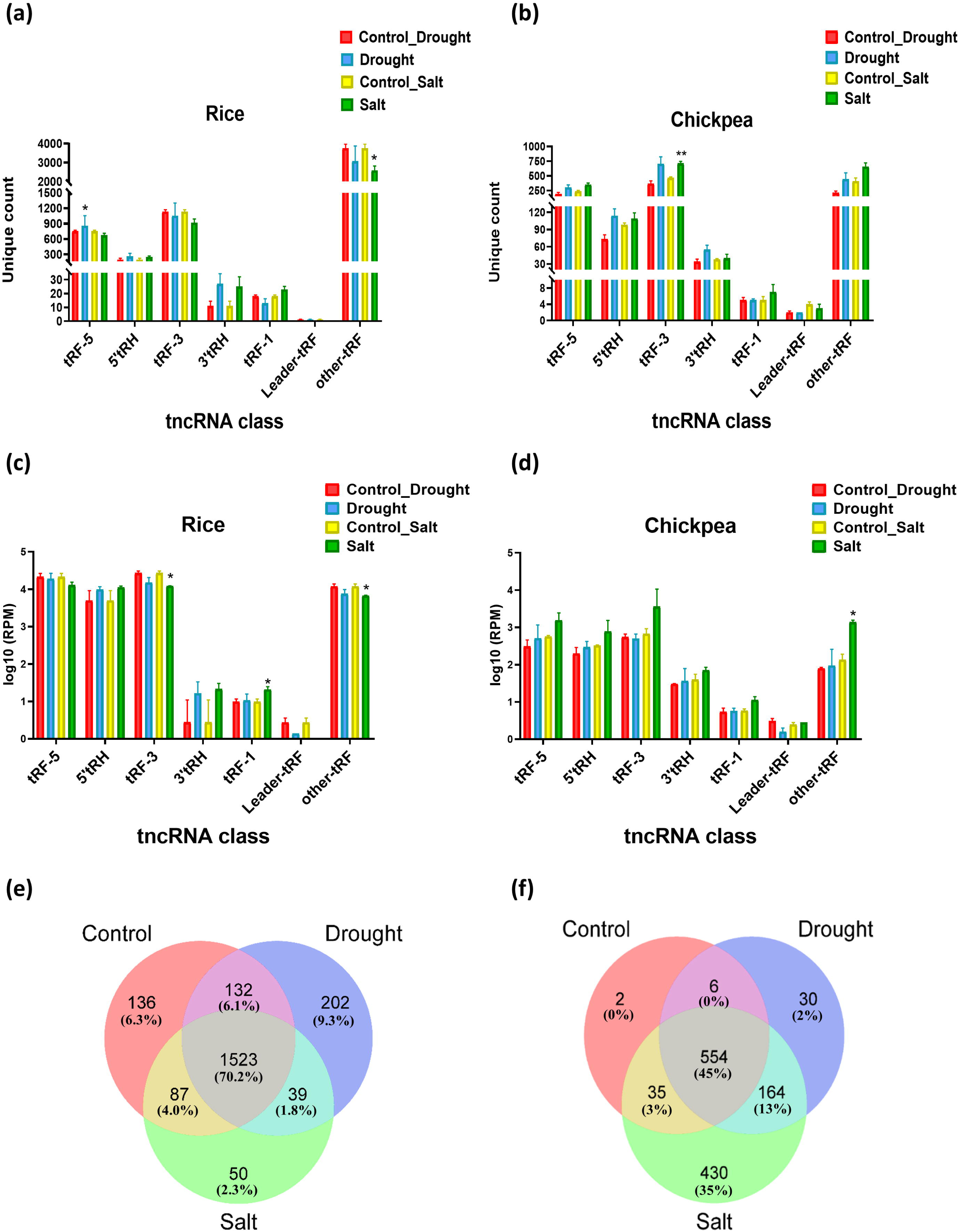
Distribution of tncRNA classes and differentially expressed tncRNAs in rice and chickpea under different conditions. (a) Category-wise distribution of unique tncRNA counts in rice and (b) Chickpea. Error bars represent the SE of three biological replicates. (c) Normalized read counts of tncRNA categories in rice and (d) Chickpea (values shown in log10 scale). Error bars represent the SE of two biological replicates. Bars marked with asterisk (*p ≤ 0.05, **p ≤ 0.01, ***p ≤ 0.001) represent statistically significant differences between control and stress conditions for each tncRNA class as determined by Student’s t-test. (e) Venn diagram illustrating the unique and shared tncRNAs in rice and (f) chickpea under control, drought, and salt stress conditions.

To comprehensively analyze the distribution and overlap of uniquely expressed tncRNAs responsive to different abiotic stresses, Venn diagrams were constructed for rice and chickpea under control, drought, and salt conditions (Figs 2e, f). In rice, a total of 1,523 tRFs (70.2%) were common across all three conditions, representing a core set of conserved tRFs, while 136 (6.3%), 202 (9.3%), and 50 (2.3%) were unique to control, drought, and salt treatments, respectively. Smaller overlaps between control–drought (132), control–salt (87), and drought–salt (39) indicated tRFs shared between specific stresses (Fig. 2e). Similarly, in chickpea (Fig. 2f), 554 tRFs (45%) were commonly expressed across all three conditions, while 2 (0%), 30 (2%), and 430 (35%) were specific to control, drought, and salt, respectively. Partial overlaps between control–drought (6), control–salt (35), and drought–salt (164) reflected distinct groups responsive to combined stress effects. Overall, the analysis highlights the complexity and specificity of tncRNA-mediated regulation under varying stress conditions, suggesting that different tncRNAs may play crucial roles in orchestrating plant responses to distinct environmental challenges.

### tncRNAs Display Differential Expression under Abiotic Stress

DESeq2 analysis of these tncRNAs under various abiotic stress conditions in rice and chickpea was performed (Supplementary Table S4). Based on the DESeq2 differential expression analysis, we selected several upregulated and downregulated tncRNAs for experimental validation using stem-loop qRT-PCR. The identified tncRNAs were grouped into two types: longer tRFs, such as tRNA halves (tRHs) longer than 25 nucleotides (usually include the anticodon loop), and shorter tRFs, mainly 5’tRFs and 3’tRFs, ranging from 19–25 nucleotides. In rice, 18 tncRNAs were validated (Table 2), and in chickpea, 31 tncRNAs were validated under abiotic stresses (Table 3).

In rice under drought stress, the 40-nucleotide 3’tRH HisGTG (tRH3002Os) and several tRF-5s (tRF5003Os, tRF5005Os, tRF5006Os, and tRF5007Os) were strongly upregulated, while tRF-1 (tRF1001Os) and tRF-3 (tRF3001Os) were significantly downregulated among the 10 validated tncRNAs (Fig. 3a). The increased levels of 3’tRHs and tRF-5s suggest that these tncRNAs may regulate pathways important for drought tolerance, such as root development and water retention. Interestingly, a dysregulation of tRF-1 was observed under stress conditions, indicating its potential but less-explored role in stress responses. Similarly, under salt stress, the 37-nucleotide 5’tRH (tRH5001Os) and several tRF-5s derived from the internal region of tRNA GluTTC (tRF5008Os, tRF5009Os, and tRFO001Os) were strongly upregulated, while other-tRF (tRFO002Os) and some tRF-5s (tRF5011Os and tRF5012Os) were downregulated among the 8 validated tncRNAs (Fig. 3b). These findings suggest that 5’tRHs and internally derived tRFs may play roles in maintaining ion balance and other protective mechanisms during salt stress.

**Figure 3:**
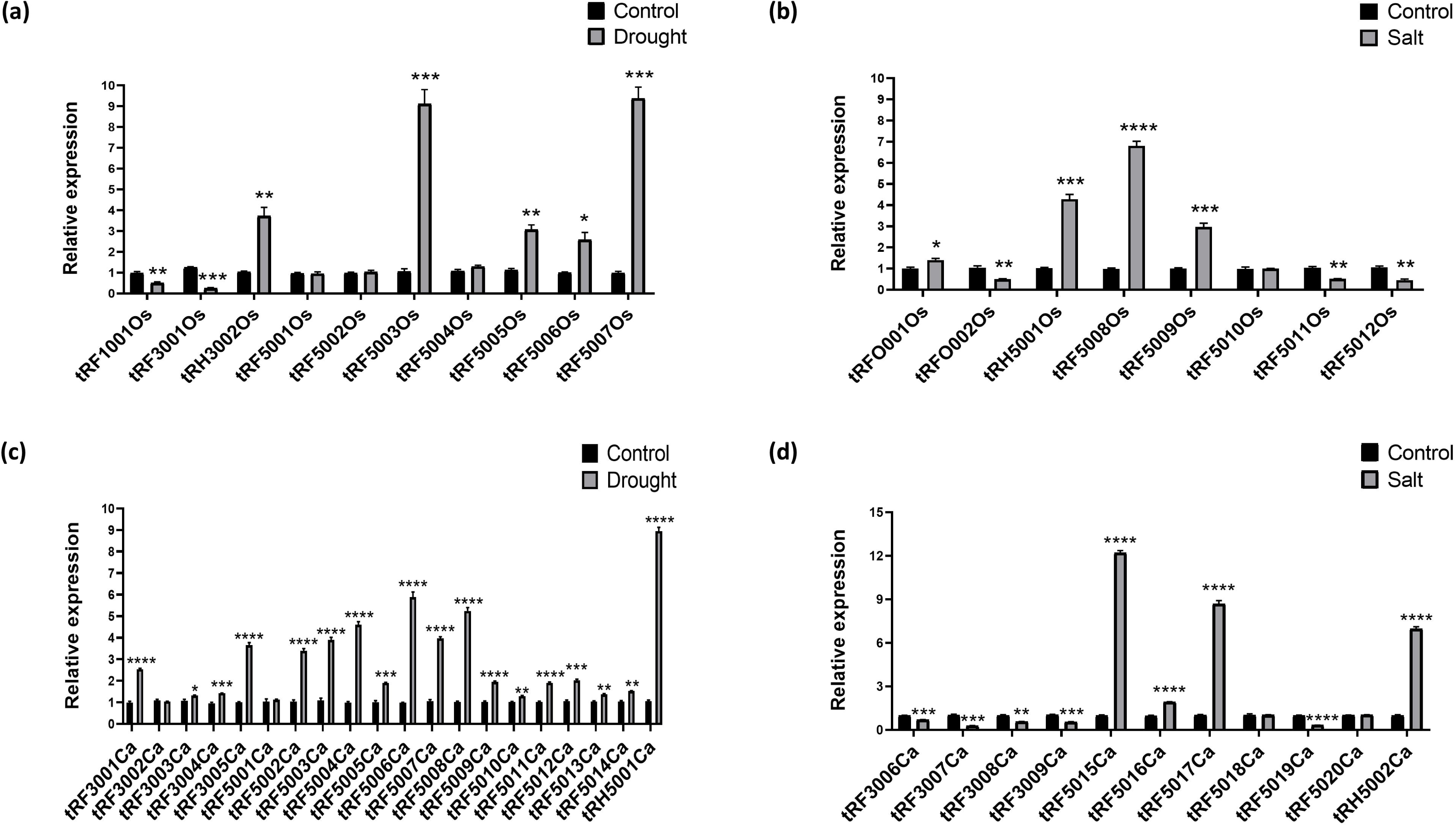
Expression of key up- and downregulated tRFs in rice and chickpea under drought and salt stress conditions. **(a)** Rice tRFs in response to drought and **(b)** salt. **(c)** Chickpea tRFs in response to drought and **(d)** salt conditions. Error bars represent the SE of three biological replicates. Bars marked with asterisk (*p ≤ 0.05, **p ≤ 0.01, ***p ≤ 0.001, ****p ≤ 0.0001) represent statistically significant differences between control and stress conditions by Student’s t-test.

In chickpea under drought stress, among the 20 validated tncRNAs, the 33-nucleotide 5’tRH (tRH5001Ca) showed the highest expression. In addition, 13 tRF-5s (tRF5002Ca, tRF5003Ca, tRF5004Ca, tRF5005Ca, tRF5006Ca, tRF5007Ca, tRF5008Ca, tRF5009Ca, tRF5010Ca, tRF5011Ca, tRF5012Ca, tRF5013Ca and tRF5014Ca) and 4 tRF-3s (tRF3001Ca, tRF3003Ca, tRF3004Ca and tRF3005Ca) were significantly upregulated, while 1 tRF-5 (tRF5001Ca) and 1 tRF-3 (tRF3002Ca) remained at levels similar to the control (Fig. 3c). Conversely, under salt stress, among the 11 validated tncRNAs, the 32-nucleotide 5’tRH (tRH5002Ca) was significantly upregulated along with 3 tRF-5s (tRF5015Ca, tRF5016Ca, and tRF5017Ca), while all tRF-3s (tRF3006Ca, tRF3007Ca, tRF3008Ca and tRF3009Ca) and 1 tRF-5 (tRF5019Ca) were significantly downregulated (Fig. 3d). The contrasting regulation under drought and salt stress suggests that specific tncRNAs have distinct roles in mediating stress responses. This highlights the complexity and specificity of tncRNA function in rice and chickpea, where differentially expressed tRFs and tRHs likely act as key post-transcriptional regulators fine-tuning gene expression for stress adaptation.

### tRF–Target Interactions Reveal Key Stress-Responsive Genes

Identification of mRNA targets is crucial for unraveling the mechanism and function of tRFs. Thus, we performed the target prediction using the psRNAtarget prediction server. We focused on shorter tRFs (19–25 nucleotides) and predicted around 80–120 targets for each in rice (Supplementary Tables S5.1, S5.2) and chickpea (Supplementary Tables S5.3, S5.4) under drought and salt stress, which showed both cleavage and translation activity. Based on these predictions, ten functionally relevant target genes linked to stress-responsive pathways were selected to construct and visualize the interaction networks between drought- and salt-associated tRFs and their corresponding targets in rice (Figs 4a, b) and chickpea (Figs 4c, d).

**Figure 4:**
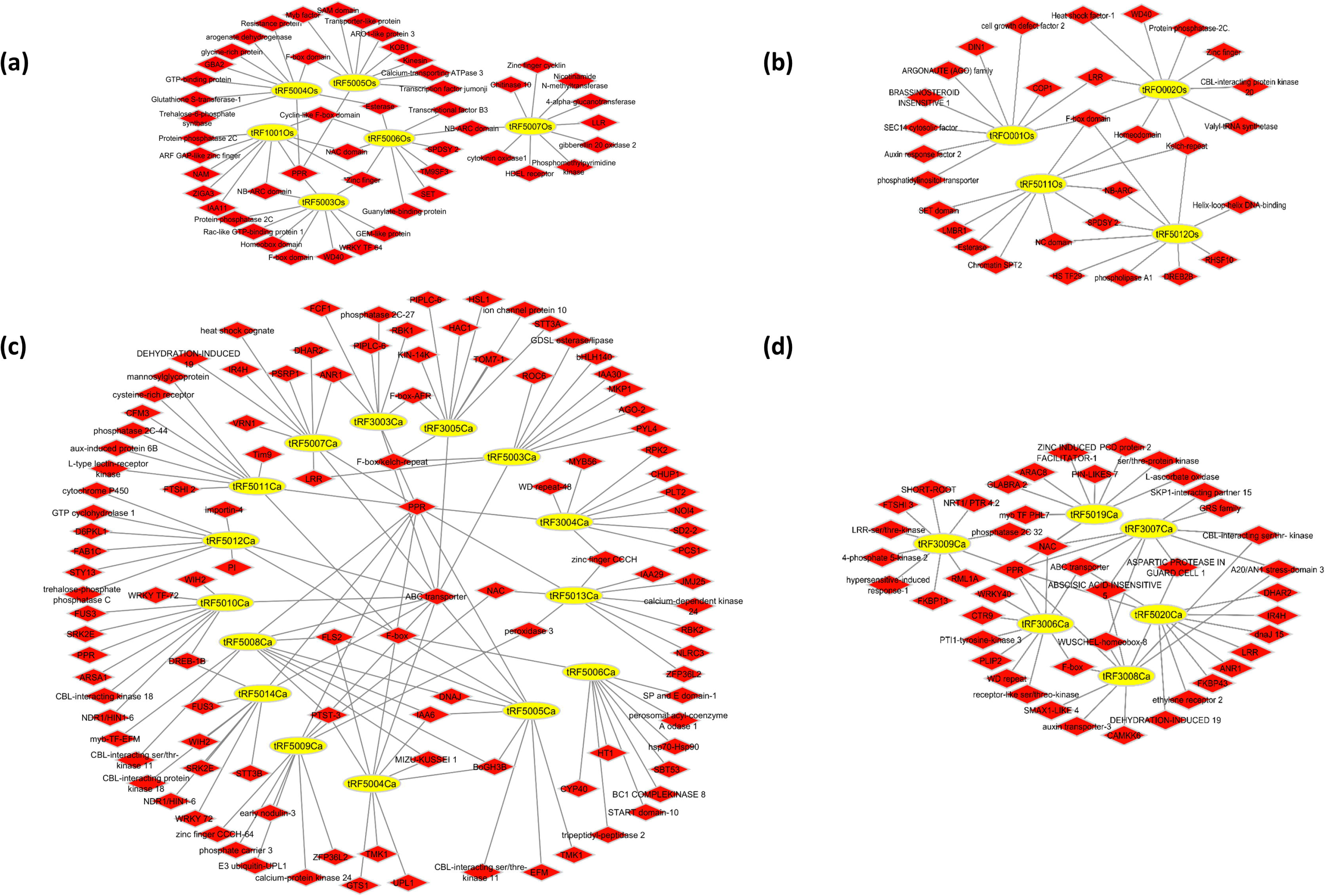
Network visualization of validated tRFs and their 10 selected stress-responsive target genes generated using Cytoscape. **(a)** Rice drought-related tRF–target interactions. **(b)** Rice salt-related tRF–target interactions. **(c)** Chickpea drought-related tRF–target interactions. **(d)** Chickpea salt-related tRF–target interactions. The networks display nodes and connections between tRFs and their annotated target genes with cleavage activity. Yellow nodes represent tRFs, while red nodes represent target genes.

Our analysis of predicted tRF targets in rice revealed that under drought stress, the 21-nt tRF-5 from ArgACG (tRF5003Os) targeted GEM-like, WRKY, and homeobox proteins, while the 25-nt tRF-5s from GlyTCC (tRF5004Os), GlyGCC (tRF5005Os), and SerTGA (tRF5007Os) were predicted to target trehalose-6-phosphate synthase, MYB, and LRR proteins, respectively. Common targets included transcripts encoding PPR, NB-ARC, NAC, F-box, and zinc finger proteins (Fig. 4a). Under salt stress, the 19-nt tRF-5 from AspGTC (tRF5012Os) targeted heat shock and DREB proteins, the 21-nt other-tRF from GluTTC (tRFO001Os) targeted auxin response factor, AGO proteins, and the 22-nt other-tRF from TyrGTA (tRFO002Os) targeted valine tRNA ligase, WD40, and CBL interacting kinase. Several domains, such as kelch repeat, F-box, homeodomain, NAC, and NB-ARC, were shared among salt-responsive tRFs (Fig. 4b). In chickpea, predicted tRF targets under drought stress showed several overlapping mRNA targets among different tRFs, including transcription factors such as NAC, WRKY, and MYB, along with ABC transporters, DREB, MIZU-KUSSEI 1, PPR, F-box, Kelch-repeat, and CBL-interacting kinase genes. The 19-nt tRF-5 from SerAGA (tRF5003Ca) targeted MKP1 and AGO2, while the 25-nt tRF-5 from GluTTC (tRF5006Ca) targeted BC1 complex kinase 8 and HSP70–HSP90 (Fig. 4c). Under salt stress, common targets across multiple tRFs included WUSCHEL-homeobox 8, F-box, CBL-interacting protein, ABC transporter, PPR, ABSCISIC ACID INSENSITIVE, and NAC. Additionally, WRKY40 and WD-repeat proteins were targeted by the 19-nt tRF-3 from CysGCA (tRF3006Ca); the 21-nt tRF-3 from ProTGG (tRF3007Ca) targeted SKP1-interacting partner 15; the 23-nt tRF-3 from ProTGG (tRF3008Ca) targeted ARR5 and SMAX1-like 4; the 25-nt tRF-3 from GlnTTG (tRF3009Ca) targeted NRT1/PTR 4.2; and the 19-nt tRF-5 from ArgCCT (tRF5019Ca) targeted MYB transcription factors (Fig. 4d). These findings suggest that tRFs may collectively regulate multiple components of stress signaling pathways in rice and chickpea. Overall, they indicate that tRFs play an important regulatory role by targeting key developmental and stress-responsive genes, thereby contributing to plant adaptation, growth, and reproduction under adverse conditions.

### Enrichment Analysis Highlights Distinct Mechanisms of Drought and Salt Response

To elucidate the biological processes, molecular functions, and pathways associated with the predicted genes, GO and KEGG pathway enrichment analyses were performed in rice (Supplementary Table S6.1) and chickpea (Supplementary Table S6.2) under drought and salt stress conditions. GO and KEGG enrichment analyses revealed distinct functional patterns of tRFs target genes in rice and chickpea under drought and salt conditions. In rice, drought-associated genes were enriched in biological processes such as plant defense response, translational elongation, meiotic chromosome condensation, and protein dephosphorylation, with roles in chloroplast inner membrane, centromeric chromosome regions, and transcription regulator complexes, and molecular functions including ADP binding and GTPase activity. KEGG analysis showed enrichment in biosynthesis of unsaturated fatty acids and sphingolipid metabolism (Supplementary Fig. S1a). Salt-associated genes in rice were significantly enriched in cellular heat acclimation, carbohydrate metabolism, and stress granule assembly, with molecular functions including transmembrane receptor kinase activity and RNA binding, as well as KEGG pathways such as glycosaminoglycan degradation and inositol phosphate metabolism (Supplementary Fig. S1c).

In chickpea, drought-associated genes were significantly enriched in RNA modification, protein catabolic processes, and protein phosphorylation, with molecular functions including ATP/ADP binding, protein kinase activity, ATP hydrolysis, and ABC transporter activity. KEGG enrichment was observed in ABC transporter pathways (Supplementary Fig. S1b). Whereas salt-associated genes were enriched in the regulation of cell shape, with the cellular component of RNA polymerase-I complex, molecular function of ribonucleoside binding, and KEGG enrichment in plant hormone signal transduction (Supplementary Fig. S1d).

Overall, the enrichment analyses suggest that drought- and salt-associated tRFs target genes in rice and chickpea are involved in distinct biological processes, molecular functions, and pathways. Drought-responsive genes are primarily linked to defense, metabolic regulation, and transporter activities, whereas salt-responsive genes are associated with cellular adaptation, signaling, and regulatory functions. These results highlight that rice and chickpea activate both shared and unique molecular mechanisms to cope with drought and salt stresses.

### RLM-RACE and qPCR Confirm Target Gene Regulation

To experimentally validate the predicted target genes, 5′ RLM-RACE was performed to confirm the tRFs cleavage sites in the corresponding target mRNAs. Three potential target genes from rice and six from chickpea, which play significant roles in various stress-related metabolic pathways, were selected for validation using 5′ RLM-RACE (Table 4). The PCR products obtained from RACE were analyzed on agarose gels (Figs 5 a, b), and the cleaved fragments were excised, cloned, and confirmed through sequencing. In rice, cleavage sites were confirmed for two tRF-5s (tRF5003Os and tRF5005Os) and one other-tRF (tRFO002Os), validating their direct targeting of mRNAs encoding an OsGEM-like protein (XM_015776041.2), a Pentatricopeptide repeat domain-containing protein (XM_015780543.2), and valine–tRNA ligase (XR_003241151.1), respectively (Fig. 5a; Table 4). tRF5003Os and tRF5005Os showed complementary regions in the 5′ UTR and 3′ UTR of their target genes, but interestingly, the cleavage sites were located upstream of these complementary regions. In contrast, tRFO002Os showed complementarity with the coding region of its target gene, and the cleavage site was detected within the predicted complementary region.

**Figure 5:**
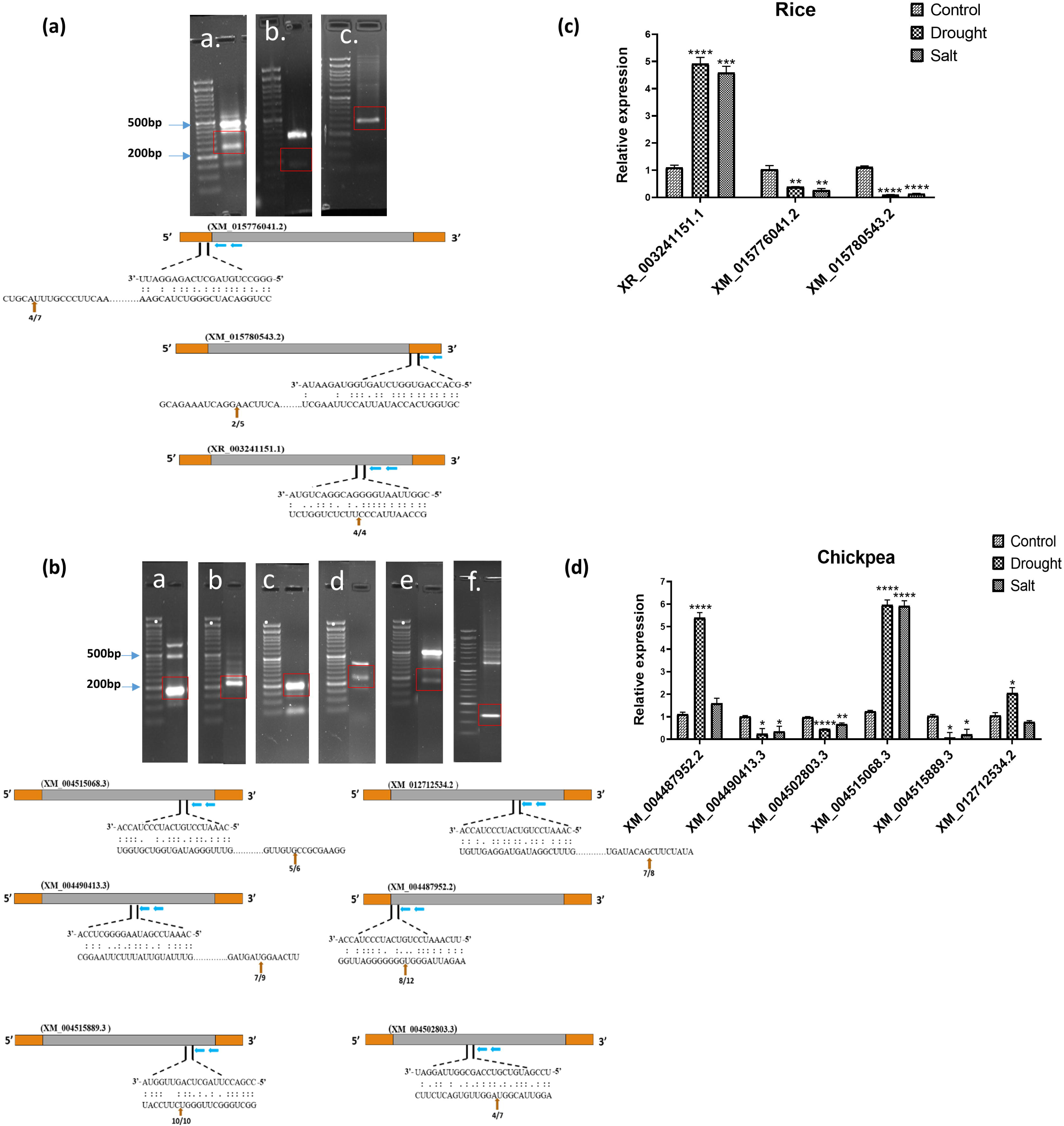
Gel electrophoresis, cleavage site validation of RLM-RACE PCR and expression profiling of target genes across different stress conditions for rice and chickpea. **(a)** Rice: lanes (a–c) show RACE products. Red boxes highlight cleaved products confirmed by sequencing. The corresponding cleavage sites within mRNA are shown below; orange boxes mark UTRs, blue arrows indicate primer positions, and red arrows indicate cleavage points by the tRFs. **(b)** Chickpea: lanes (a–f) show cleavage products of mRNA targets. **(c)** Relative expression levels of rice and **(d)** chickpea target genes in drought and salt stress conditions compared to the control. Error bars indicate the SE of three biological replicates. Bars marked with asterisk (*p ≤ 0.05, **p ≤ 0.01, ***p ≤ 0.001, ****p ≤ 0.0001) represent statistically significant differences between control and stress conditions by student’s t-test.

**Table 4:**
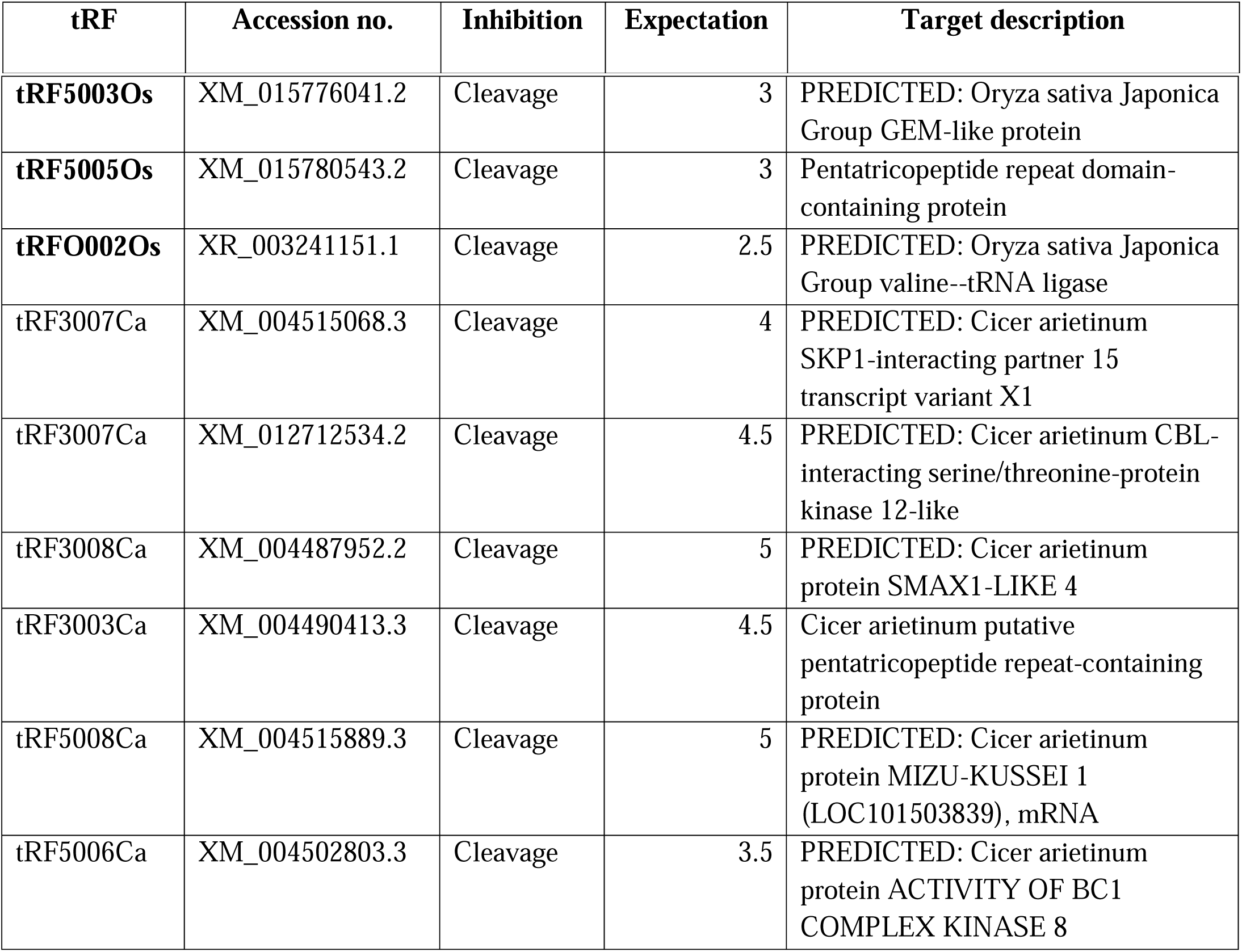
List of mRNA targets of the differentially expressed target transcripts used for RLM-RACE in Rice (Figure 6a) and Chickpea (Figure 6b).

| tRF | Accession no. | Inhibition | Expectation | Target description |
| --- | --- | --- | --- | --- |
| tRF5003Os | XM_015776041.2 | Cleavage | 3 | PREDICTED: Oryza sativa Japonica Group GEM-like protein |
| tRF5005Os | XM_015780543.2 | Cleavage | 3 | Pentatricopeptide repeat domain-containing protein |
| tRFO002Os | XR_003241151.1 | Cleavage | 2.5 | PREDICTED: Oryza sativa Japonica Group valine--tRNA ligase |
| tRF3007Ca | XM_004515068.3 | Cleavage | 4 | PREDICTED: Cicer arietinum SKP1-interacting partner 15 transcript variant X1 |
| tRF3007Ca | XM_012712534.2 | Cleavage | 4.5 | PREDICTED: Cicer arietinum CBL-interacting serine/threonine-protein kinase 12-like |
| tRF3008Ca | XM_004487952.2 | Cleavage | 5 | PREDICTED: Cicer arietinum protein SMAX1-LIKE 4 |
| tRF3003Ca | XM_004490413.3 | Cleavage | 4.5 | Cicer arietinum putative pentatricopeptide repeat-containing protein |
| tRF5008Ca | XM_004515889.3 | Cleavage | 5 | PREDICTED: Cicer arietinum protein MIZU-KUSSEI 1 (LOC101503839), mRNA |
| tRF5006Ca | XM_004502803.3 | Cleavage | 3.5 | PREDICTED: Cicer arietinum protein ACTIVITY OF BC1 COMPLEX KINASE 8 |

In chickpea, cleavage sites were confirmed for three tRF-3s (tRF3003Ca, tRF3007Ca, and tRF3008Ca) and two tRF-5s (tRF5006Ca and tRF5008Ca). For tRF3007Ca, two predicted target cleavage sites were validated in mRNAs encoding SKP1-interacting partner 15 (XM_004515068.3) and CBL-interacting serine/threonine-protein kinase 12-like (XM_012712534.2), while tRF3003Ca targeted a pentatricopeptide repeat domain-containing protein (XM_004490413.3). Both tRFs had complementary regions in the coding sequence, with cleavage sites located downstream of the complementary regions (Fig. 5b; Table 4). In contrast, tRF3008Ca cleaved SMAX1-LIKE 4 protein (XM_004487952.2), tRF5006Ca cleaved ACTIVITY OF BC1 COMPLEX KINASE 8 protein (XM_004502803.3), and tRF5008Ca cleaved MIZU-KUSSEI 1 (XM_004515889.3), with cleavage sites detected within the complementary regions at different positions.

Further qPCR analysis of RLM-RACE validated target genes revealed a negative correlation between tRFs and their target mRNAs in both rice and chickpea (Figs 5c, d). In rice, highly induced tRFs under drought and salt stress were associated with significant downregulation of their target genes, OsGEM-like protein (XM_015776041.2) and Pentatricopeptide repeat domain-containing protein (XM_015780543.2). In contrast, valine–tRNA ligase (XR_003241151.1), which corresponded to lower-abundance tRFs, was upregulated under both stress conditions (Fig. 5c). Similarly, in chickpea, upregulated tRFs led to significant downregulation of target genes, including Pentatricopeptide repeat protein (XM_004490413.3), BC1 COMPLEX KINASE 8 protein (XM_004502803.3), and MIZU-KUSSEI 1 (XM_004515889.3). Conversely, target genes of downregulated tRFs, including SKP1-interacting partner 15 (XM_004515068.3), CBL-interacting serine/threonine-protein kinase 12-like (XM_012712534.2), and SMAX1-LIKE 4 proteins (XM_004487952.2), were significantly upregulated (Fig. 5d). Collectively, the results indicate that tRFs mediate site-specific cleavage of their target mRNAs, thereby modulating gene expression. Importantly, this cleavage does not solely lead to transcript degradation; in certain cases, it may also facilitate upregulation of the target genes, indicating a potential feedback role for tRFs in fine-tuning post-transcriptional regulation.

### Subcellular localization of target genes

To study the subcellular localization of selected genes, *Agrobacterium tumefaciens* GV3101 carrying 35S:GFP (control), 35S:*OsGEM*-GFP, 35S:*OsPPR*-GFP, 35S:*CaSKP1*-GFP, and 35S:*CaMIZ1*-GFP constructs (schematic shown in Supplementary Fig. S3) was co-infiltrated into *Nicotiana benthamiana* leaves along with plasma membrane (PM-rk-mCherry) or nuclear (NM-rk-mCherry) markers. The control 35S:GFP signal was uniformly distributed throughout the cell (Supplementary Fig. S4). In contrast, the OsGEM-GFP fusion protein showed strong nuclear localization (Fig. 6a), while OsPPR-GFP was observed at both the plasma membrane and nuclear envelope, indicating dual localization (Fig. 6b). The CaSKP1-GFP and CaMIZ1-GFP fusion proteins predominantly localized to the plasma membrane (Figs 6c, d).

**Figure 6:**
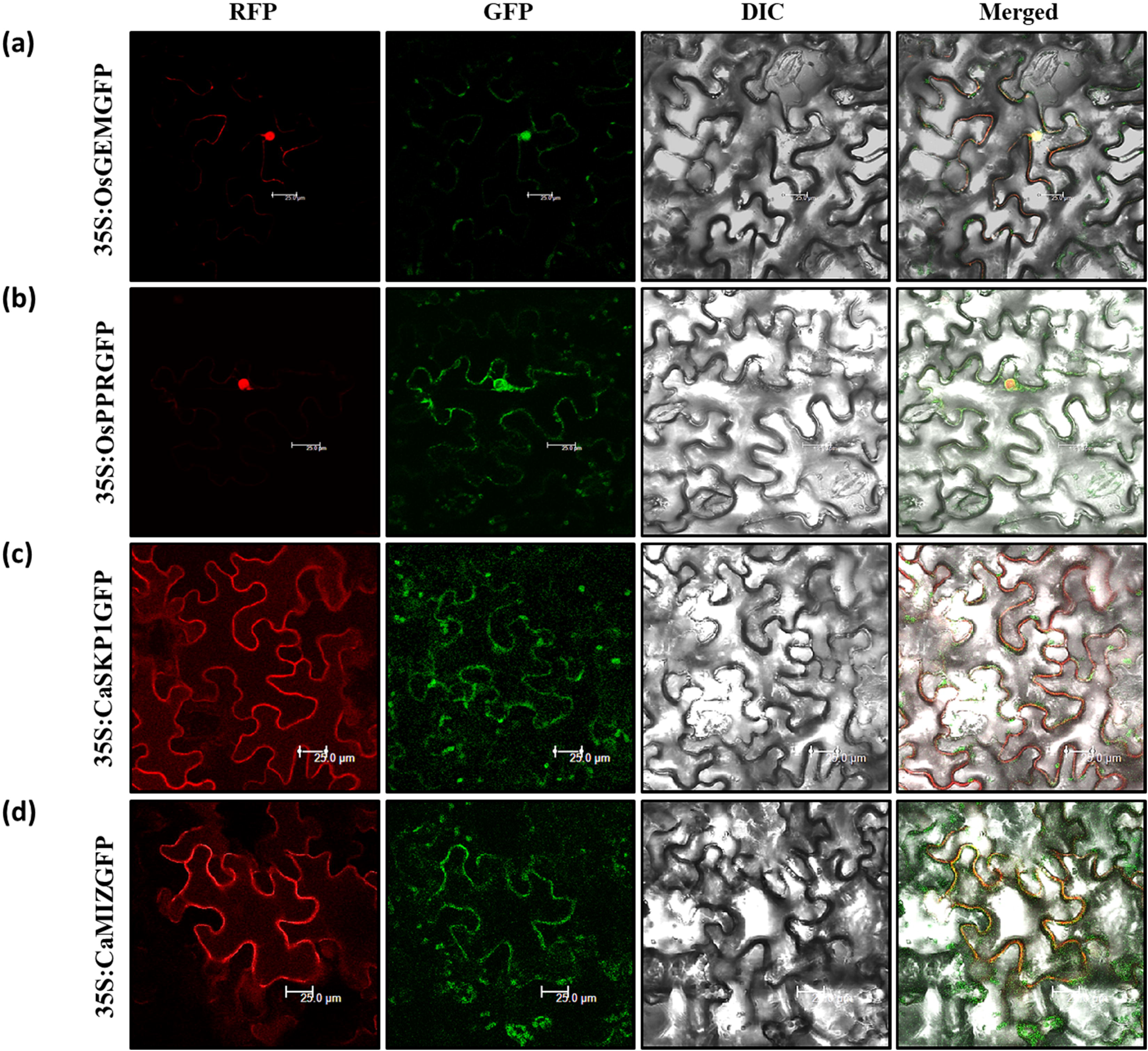
Subcellular localization of target genes of rice and chickpea in tobacco leaf epidermal cells. **(a)** Colocalization of 35S:OsGEMGFP and **(b)** 35S:OsPPRGFP with NM-rk-mCherry (Nuclear Marker). **(c)** Colocalization of 35S:CaSKP1GFP and **(d)** 35S:CaMIZ1GFP with PM-rk-mCherry (Plasma membrane Marker). Confocal microscopy was used to examine the GFP expression after 48 hours of co-infiltration in N. benthamiana leaves. Scale bar: 25µm.

## Discussion

We identified a diverse range of stress-responsive tncRNAs in rice and chickpea, revealing both conserved and species-specific expression patterns under drought and salinity stress. These findings reinforce that tRFs are produced through regulated processes rather than random tRNA degradation, consistent with earlier reports (Kumar et al., 2014; Alves et al., 2017), supporting the idea that tRFs/tncRNAs are not simply by-products of random tRNA degradation but rather arise through regulated mechanisms. Furthermore, the findings highlight that both tRF length and the choice of parental tRNA show species-specific patterns under specific conditions (Table 2, 3).

In rice, drought stress led to a significant upregulation of the 3′tRH derived from HisGTG along with several tRF-5s derived from ArgACG, GlyGCC, AspGTC, and SerTGA (Table 2; Fig. 3a). Interestingly, under salt stress, the expression of 5′tRHs, including tRF-5s derived from the same anticodon GluTTC, as well as other-tRFs, was also markedly upregulated (Table 2; Fig. 3b). This observation aligns with recent research reporting the significant induction of tRF-5 derived from ArgTCG in rice under oxidative stress (Huang et al., 2025). While in chickpea, tRF-5s derived from GluTTC, SerAGA, TyrGTA, GluTTC, MetCAT, GluCTC, GlnTTG, AlaCGC, AlaTGC and GluCTC were found to be upregulated, whereas tRF-3s derived from the same anticodon ThrGGT displayed both upregulation and no significant change under drought stress (Table 3; Fig. 3c). Similarly, under salt stress, tRF-5s derived from the same anticodon GluTTC also showed variable expression patterns, with some being upregulated while others remained unchanged (Table 3; Fig. 3d), highlighting their specific role in regulation. These stress-specific expression patterns suggest that tncRNAs may function in distinct adaptive roles, contributing to drought tolerance on one hand, and to maintaining ion balance and activating defence mechanisms under salt stress on the other. Previous research has shown that certain tncRNAs are specifically responsive to different abiotic stresses. For example, in *Arabidopsis*, 5′ tRF ArgCCT is elevated under both drought and oxidative stress (Alves et al., 2017), while 5′ tRF GlyGCC, initially induced by salt stress, also responds to UV exposure (Cognat et al., 2017). Similarly, in wheat seedlings, the 3′ tRF TyrGUA has been reported to react to heat, salt, and drought conditions (Wang et al., 2016). Notably, the drought-induced 5′ tRF ArgCCT observed in *Arabidopsis* does not exhibit the same response in rice (Alves et al., 2017), suggesting that the regulation of identical tncRNAs can differ across plant species.

Identifying the mRNA targets of tRFs is essential for deciphering their regulatory functions and underlying mechanisms. Target prediction using the psRNATarget server revealed that Pentatricopeptide repeat (PPR) proteins represent prominent targets, particularly for tRF-5s derived from ArgACG and GlyGCC, as well as tRF-3s derived from GlyGCC, ProTGG, and GlnTTG under different stress conditions in rice and chickpea (Supplementary Table S5). The predicted interactions were further validated by RLM-RACE, which confirmed that the cleavage site of the PPR transcript was located upstream of the complementary region in rice and downstream in chickpea (Figs 5a, b). Finally, qPCR analysis verified these findings, showing downregulation of PPR expression following cleavage (Figs 5c, d), highlighting a direct regulatory role of tRFs in modulating PPR expression under stress. PPR proteins are sequence-specific RNA-binding factors that regulate post-transcriptional processes in mitochondria and chloroplasts, including RNA editing, splicing, stability, and translation (Barkan and Small, 2014; Schmitz-Linneweber and Small, 2008). Several PPRs are linked to abiotic stress responses; for instance, *OsPPR674* influences mitochondrial RNA editing and drought tolerance in rice (Li et al., 2025), while *SOAR1* regulates ABA signaling and enhances stress tolerance in *Arabidopsis* (Mei et al., 2014). Previous findings indicate that PPR proteins contribute to both growth and stress adaptation in crops, and our identification of stress-responsive tncRNAs potentially targeting PPR transcripts in rice and chickpea suggests a plausible mechanism for species-specific modulation of organelle gene expression during stress adaptation. Similarly, under drought stress, the tRF-5 derived from ArgACG was found to target the mRNA encoding GEM-like protein 1 (Fig. 4a), with the cleavage site located upstream of the complementary region in rice (Fig. 5a). qPCR analysis confirmed that its expression was downregulated (Fig. 5b). Since GEM-like proteins have been associated with developmental processes (seed development and inflorescence architecture) and the ABA signaling pathway (Baron et al., 2014; Mauri et al., 2016), this suggests that the tRF may help regulate key developmental pathways to support plant growth and reproduction under stress conditions.

Interestingly, in chickpea, the hydrotropism-related gene *MIZU-KUSSEI 1* (*MIZ1*), which promotes drought avoidance by directing root growth toward water (Kobayashi et al., 2007; Shkolnik et al., 2018; Yamazaki et al., 2012), was identified as a target of a tRF-5 derived from TyrGTA. This aligns with findings in *Arabidopsis*, where overexpression and knockdown of 5′ tRF-Ala affected plant growth and arsenite sensitivity during root development (Li et al., 2023), highlighting the regulatory role of tRFs in root growth and stress adaptation. Additionally, in chickpea, a tRF-5 derived from GluTTC was found to target *BC1 Complex Kinase 8* (*ABC1K8*) (Fig. 4c). Both of these interactions were validated through cleavage site detection within the complementary region (Fig. 5b), resulting in reduced transcript expression (Fig. 5d). ABC1K8, a member of the atypical kinase family, is known to regulate the production of reactive oxygen species (ROS), thereby modulating plant responses to oxidative stress (Lundquist et al., 2013; Manara et al., 2014; Martinis et al., 2014). In contrast, tRF-3s derived from ProTGG were predicted to target and cleave CBL-interacting serine/threonine-protein kinase 12 (Fig. 5c), a key component of calcium-mediated signaling in abiotic stress responses (Batistič and Kudla, 2009; Cheong et al., 2010; Du et al., 2021), SKP1-interacting partner 15 (Fig. 5c), a part of the SCF ubiquitin ligase complex regulating protein degradation and hormone signaling (Li et al., 2012; Varshney et al., 2023; Zhao et al., 2003), and SMAX1-like 4, a regulator of karrikin and strigolactone signaling involved in seedling growth and stress adaptation (Feng et al., 2023; Stanga et al., 2013; Yang et al., 2020). Intriguingly, qPCR analysis revealed upregulation of these genes despite cleavage (Fig. 5d), suggesting that partial degradation by tRFs does not necessarily reduce transcript abundance and may, in some cases, contribute to transcript upregulation (Kumar et al., 2016; Sobala and Hutvagner, 2011). Taken together, these findings suggest that tRFs appear to fine-tune multiple stress-responsive mechanisms rather than simply silencing genes.

The enrichment patterns indicated that these targets participate in diverse biological processes, molecular functions, and pathways in both rice and chickpea (Supplementary Fig. S1). Notably, genes responsive to drought stress were predominantly associated with defense responses, metabolic regulation, and transporter activity (Muthuramalingam et al., 2017; Shinozaki and Yamaguchi-Shinozaki, 2007), whereas those responsive to salt stress were enriched in categories linked to cellular adaptation, signaling, and regulatory mechanisms (Deinlein et al., 2014; Zhu, 2016). These observations suggest that while both crops engage shared and unique stress-related pathways, they also employ crop-specific molecular strategies to withstand drought and salinity challenges, consistent with earlier reports highlighting both conserved and species-specific mechanisms of stress adaptation in cereals and legumes (Fang and Xiong, 2015; Varshney et al., 2011).

Additionally, subcellular localization analyses were conducted on the target genes in both rice and chickpea. In rice, OsGEM was predominantly localized in the nucleus (Fig. 6a), while OsPPR was detected in both the plasma membrane and nuclear envelope (Fig. 6b). These findings align with previous studies reporting the nuclear localization of GRAM protein (OsABAR1) in rice (Zheng et al., 2020) and the dual localization of PPR in *Arabidopsis* (Colcombet et al., 2013). In chickpea, CaSKP1 and CaMIZ1 were primarily localized to the plasma membrane, with light patches observed in the cytosol (Figs 6c, d). This localization pattern is consistent with earlier findings in *Arabidopsis*, where ASK20A–GFP and ASK20B–GFP signals were detected in the cytosol (Ogura et al., 2008), and AtMIZ1 was shown to associate with the cytoplasmic side of the endoplasmic reticulum (Yamazaki et al., 2012), suggesting their involvement in signal perception and transduction at the cellular periphery during stress responses. Collectively, these spatial distributions, together with the regulatory effects of tRFs on their transcripts, indicate that target proteins may coordinate stress adaptation both at the transcriptional level and through compartment-specific functions, highlighting the multi-layered role of tRF-mediated regulation in plant stress tolerance.

## Conclusion

We identified and characterized novel tRNA-derived fragments (tRFs) in rice and chickpea that regulate a diverse set of stress-responsive genes under drought and salinity conditions. Our analyses demonstrated that tRFs target key regulators involved in root hydrotropism, ROS homeostasis, hormone signaling, protein degradation, and developmental processes. Interestingly, while some target transcripts were downregulated following cleavage, others were upregulated, suggesting that tRFs act not only as negative regulators but also as fine-tuners of gene expression, contributing to dynamic stress adaptation. Overall, our findings underscore the multilayered regulatory potential of tRFs in plant stress responses, as illustrated in a schematic model (Fig. 7). Future studies using approaches such as tRF overexpression through artificial miRNA (amiRNA) technology and short tandem target mimic (STTM) systems will be valuable for uncovering their mechanistic roles in stress regulation. Moreover, CRISPR-Cas–based genome editing can be employed to modify key genes involved in tRF biogenesis, such as *Dicer-like* or *RNase Z,* and their interacting partners, to better understand their regulatory networks. Together, these strategies could provide deeper functional insights and open new possibilities for developing crops with improved tolerance to abiotic stresses.

**Figure 7:**
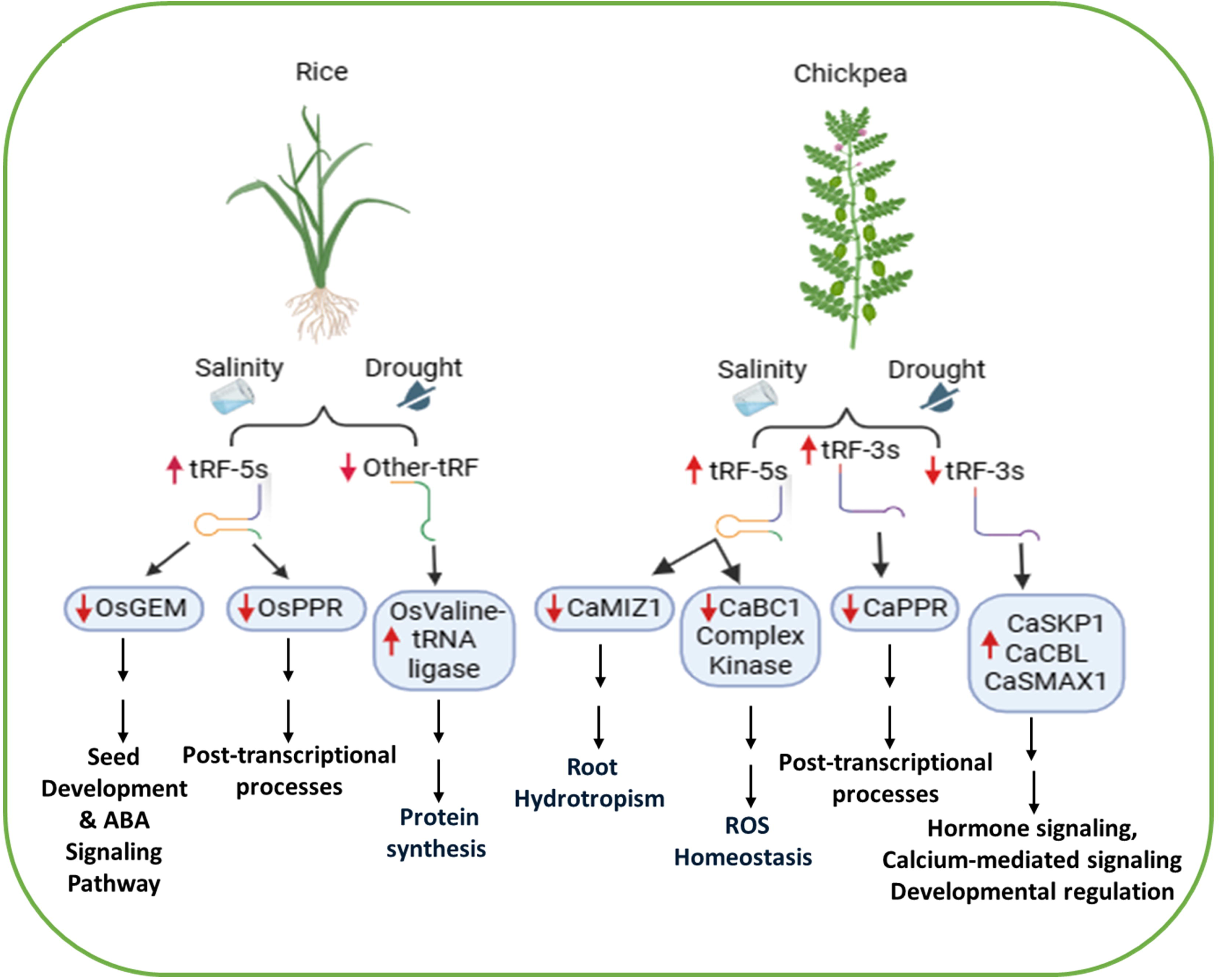
A proposed model illustrating the diverse roles of tRFs in regulating gene expression under stress responses in rice and chickpea. Stress conditions promote tncRNA production, with tRF-5 and tRF-3 acting as key regulators of stress-responsive genes. Elevated tncRNA levels suppress the expression of their target genes, while reduced tRF expression results in gene upregulation. Specifically, tRF-5s target OsGEM, OsPPR, CaMIZ1, and CaBC1, leading to repression of genes associated with seed development, ABA signaling, post-transcriptional regulation, root hydrotropism, and ROS homeostasis. In contrast, other-tRF and tRF-3s target OsValine-tRNA ligase, CaSKP1, CaCBL1, and CaSMAX1, thereby enhancing the expression of genes involved in protein synthesis, hormone signaling, calcium-mediated signaling, and developmental regulation. Red upward arrows denote upregulation, while downward arrows denote downregulation.

## Data Availability

Small RNA sequencing data supporting this study are available at the NCBI Sequence Read Archive (SRA) under accession number PRJNA1362969. All pipeline scripts and codes for tncRNA prediction are freely available at http://nipgr.ac.in/tncRNA, and https://github.com/skbinfo/tncRNA-Toolkit.

## Competing interests

The authors have no conflicts of interest to declare.

## Funding

This research is supported by the BT/PR40146/BTIS/137/4/2020 project grants by the Department of Biotechnology (DBT), Government of India, and by the core grant of the National Institute of Plant Genome Research (NIPGR) in the laboratory of SK.

## Author contributions

SZ: RNA extraction of chickpea, Insilco identification of tncRNAs, Experimental validation of tncRNAs, RLM-RACE, Data curation, Formal analysis, and Writing-Original draft; RG: Experimental validation of tncRNAs and RLM-RACE, Data curation, Formal analysis, and Writing-Original draft; ST: Subcellular localization experiments, Data curation, Formal analysis, and Writing-Original draft; DKB: RNA extraction of rice; AS: Discussions; SK: Conceptualization; Funding acquisition, Project administration, Resources, Supervision, and Writing-review & editing.

## Acknowledgments

The authors are thankful to the Department of Biotechnology (DBT)-eLibrary Consortium, India, for providing access to e-resources. The authors acknowledge the Computational Biology & Bioinformatics Facility (CBBF) of the National Institute of Plant Genome Research (NIPGR).

## Notes

### Competing Interest Statement

The authors have declared no competing interest.

